# The Medical Genome Reference Bank: a whole-genome data resource of 4,000 healthy elderly individuals. Rationale and cohort design

**DOI:** 10.1101/274019

**Authors:** Paul Lacaze, Mark Pinese, Warren Kaplan, Andrew Stone, Marie-Jo Brion, Robyn L Woods, Martin McNamara, John J McNeil, Marcel E Dinger, David M Thomas

## Abstract

Allele frequency data from human reference populations is of increasing value for filtering and assignment of pathogenicity to genetic variants. Aged and healthy populations are more likely to be selectively depleted of pathogenic alleles, and therefore particularly suitable as a reference populations for the major diseases of clinical and public health importance. However, reference studies of the healthy elderly have remained under-represented in human genetics. We have developed the Medical Genome Reference Bank (MGRB), a large-scale comprehensive whole-genome dataset of confirmed healthy elderly individuals, to provide a publicly accessible resource for health-related research, and for clinical genetics. It also represents a useful resource for studying the genetics of healthy aging. The MGRB comprises 4,000 healthy, older individuals with no reported history of cancer, cardiovascular disease or dementia, recruited from two Australian community-based cohorts. DNA derived from blood samples will be subject to whole genome sequencing. The MGRB will measure genome-wide genetic variation in 4,000 individuals, mostly of European decent, aged 60-95 years (mean age ≥ 75 years). The MGRB has committed to a policy of data sharing, employing a hierarchical data management system to maintain participant privacy and confidentiality, whilst maximizing research and clinical usage of the database. The MGRB will represent a dataset of international significance, broadly accessible to the clinical and genetic research community.

## Introduction

Each individual differs from another at millions of sites in the genome. One of the key challenges in the interpretation of whole genome sequencing (WGS) data for the diagnosis of inherited disease is discriminating rare candidate disease-causing variants from the large numbers of benign variants unique to each individual. Reference populations are powerful filters to distinguish pathogenic from population-based genetic variation; both clinically for Mendelian disorders [1,2], and in research for studies of genetic disease [3].

The availability of population-based allele frequency data has been instrumental in enabling variant filtering, assignment of pathogenicity, and frequency-based estimates of penetrance in recent years [4–6]. Variant frequency data has facilitated the diagnosis and discovery of an unprecedented number of disease-driving pathogenic mutations, and allele frequency-based filtering has become a mainstay of clinical genetics. This initially was made possible by public access to the International HapMap [6] and 1000 Genomes [5] datasets, and then more recently by the Exome Aggregation Consortium (ExAC) [4]. All of these reference projects have been pivotal in influence on human genetics, primarily due to the common aspect of data sharing. An increasing number of reference population sequencing projects are now underway worldwide, reflecting the need to understand the underlying genetic variation in different backgrounds, especially those non-European [7–11].

Despite the value provided by these genetic reference datasets, each has been limited in some way. Perhaps the most common and significant limitation has been the lack of detailed phenotypic or clinical information, at least made available to the public. Such information is of particular importance for confirming or excluding genetic disease phenotypes. Access to longitudinal clinical outcome data to interpret genetic variation considered to be pathogenic has also been lacking. For example, a cancer-free individual sampled at age 45 may go on to develop cancer at a later age. Such individuals in reference datasets may carry pathogenic variants, but still be used as negative controls in many subsequent data analyses. When combined with the stochastic and environmentally dependent nature of disease phenotypes, identification of genetically risk-deplete controls is critical to understanding the genetic basis of common diseases.

Only one whole-genome sequenced population comprised of individuals confirmed to be deplete of genetic disease phenotypes has been generated to date [12]. Depletion of disease phenotypes should decrease the burden of penetrant pathogenic mutations. Such populations, such as the one we describe here, have increased power to act as negative controls for variant filtering and assignment of pathogenicity in studies focused on inherited disease.

Another challenge of reference data sets is the size of the cohort. The larger a reference population is, the more likely the population will be to contain rare variants. Therefore, larger sample sizes provide more accurate and reliable population allele frequencies, especially at the rare end of the spectrum. This is of critical importance given that most pathogenic alleles are rare with a minor allele frequency (MAF) <0.01, particularly those of high penetrance (often MAF<0.001), and also because the majority of all single-nucleotide variants (SNVs) are rare, as shown by recent population-based whole-exome and -genome sequencing efforts, whereby singletons were by far the most abundant SNV frequency class detected [4,11,13].

Another limitation is that many human genetic reference populations have comprised of varying ethnicities, ages and genetic backgrounds, often taken from various cohorts or case-control studies. This aggregation approach helps reach higher sample numbers, but does not ensure consistency of phenotype, or confidence in the absence of disease. Achieving the unique combination of all features required for the optimal reference data set is challenging, but should include: large size by sample number, whole-genome coverage, ability to detect complex and structural variation, confirmation of health and age phenotypes, public data access to both genomic and phenotypic data, measurement of genetic sequence variation using a consistent and compatible sequencing technology (see Table 1).

**Table 1.**
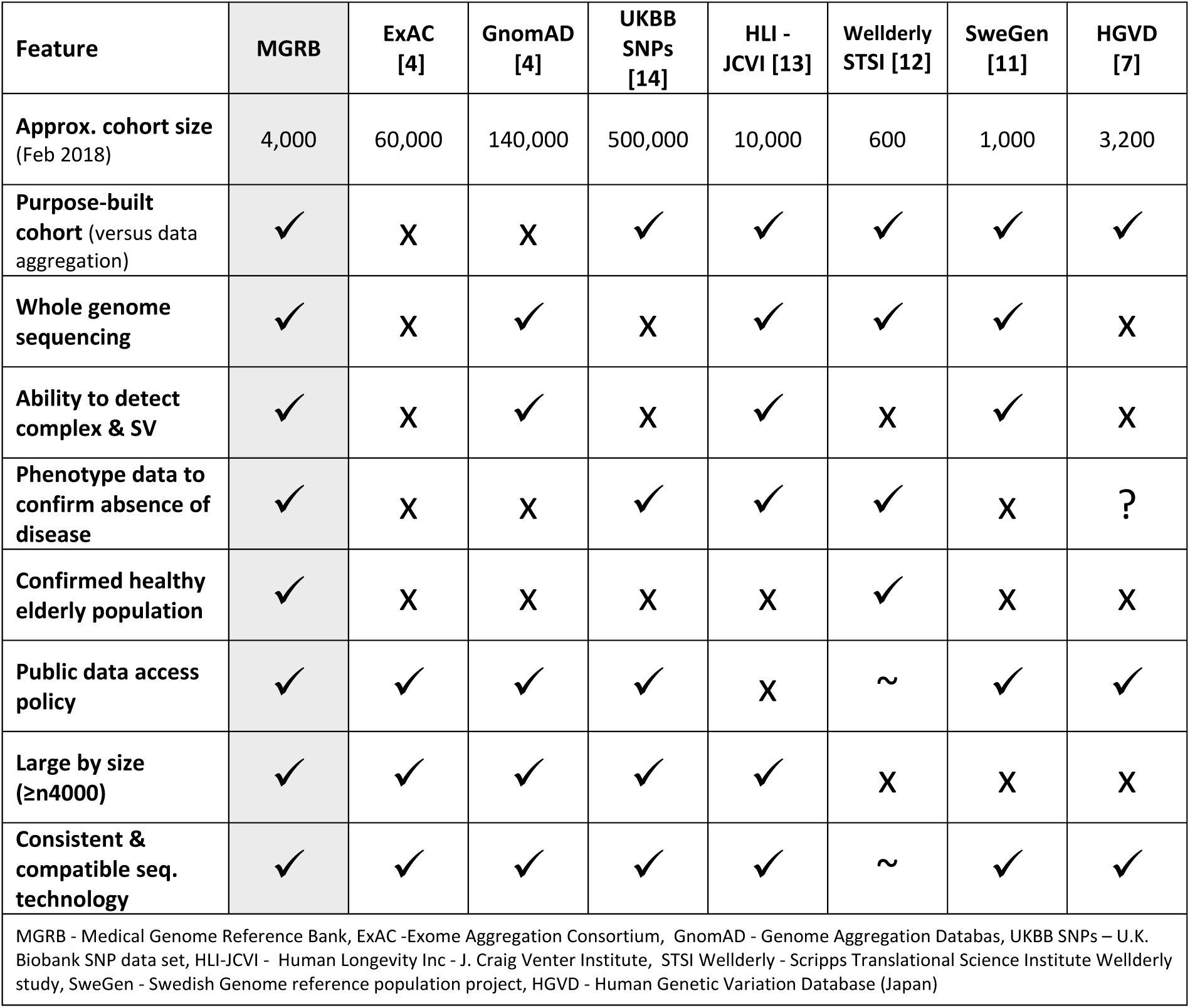
Features of human genetic reference populations (as publicly stated February 2018)

Here we present the rationale and design of the first human reference population comprised of thousands of whole genomes from confirmed healthy elderly individuals, depleted of common and rare genetic disease phenotypes. Samples for this project have been provided from two leading Australian community-based cohort studies, with access to phenotypic and clinical information. This information has been used to confirm the absence of cardiovascular disease, dementia and cancer in all participants.

The Medical Genome Reference Bank (MGRB) will involve whole genome sequencing of 4,000 healthy older adults. These individuals are participants of the ASPirin in Reducing Events in the Elderly (ASPREE) study, an international clinical trial for daily low-dose aspirin use in older people coordinated by the Department of Epidemiology & Preventive Medicine at Monash University Monash University [15], and The 45 and Up study, the largest ongoing study of healthy aging in the Southern Hemisphere, coordinated by the Sax Institute [16]. Each MGRB sample will be sequenced using Illumina technology at a minimum 30x coverage. Data will be processed using WGS best practice pipelines (GATK-BWA) and resulting population allele frequency data will be made openly-accessible and downloadable via public website. Individual-level variant-call files (VCFs), core phenotypes, and access to alignment files (BAMs) will be open to application via the MGRB Data Access Committee. Access to additional clinical information and phenotype data will be available via application to contributing cohorts, via existing data access and governance arrangements. For MGRB project overview, see Figure 1.

**Figure 1:**
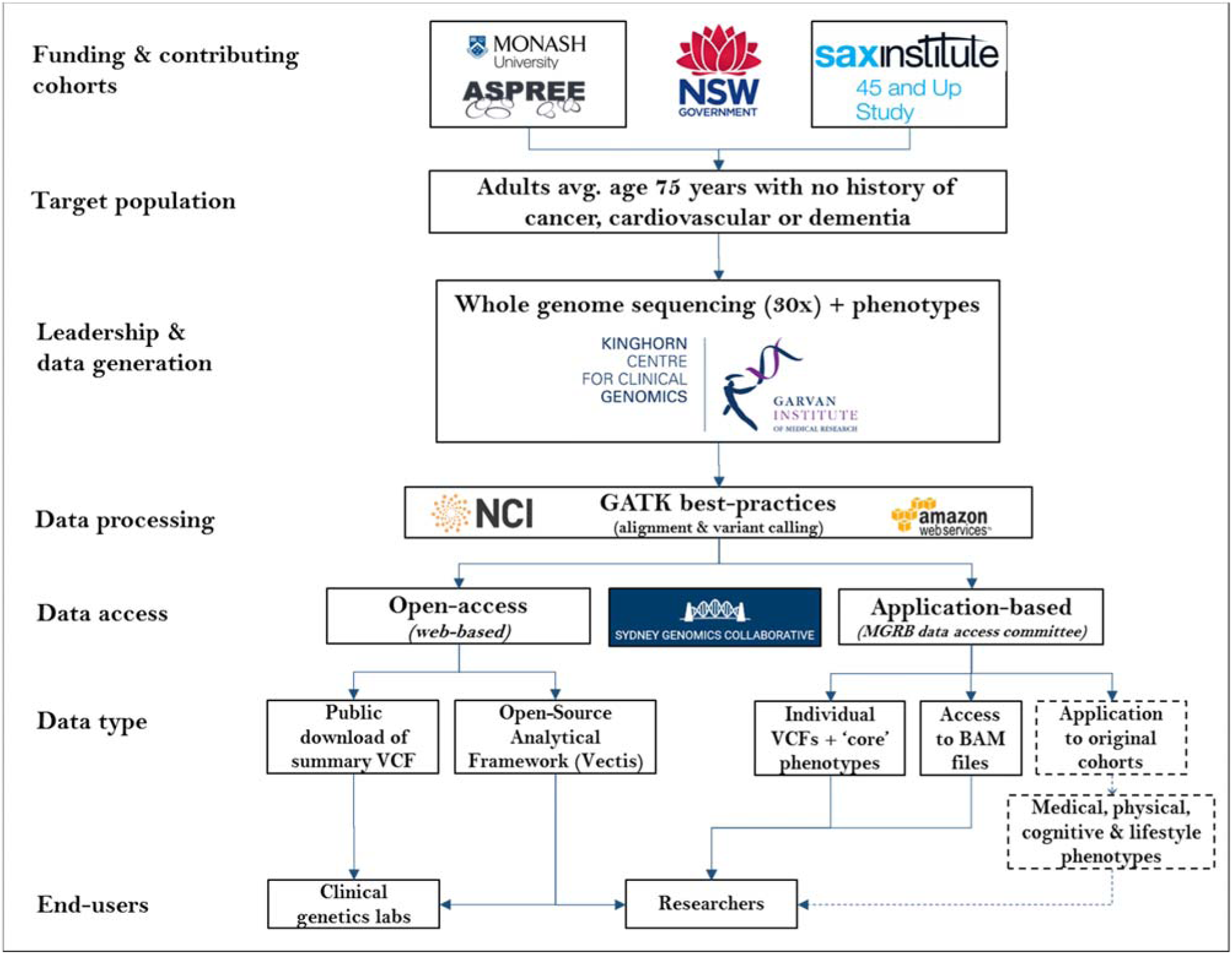
The Medical Genome Reference Bank: Project overview.

## Methods/Design

### Inclusion/Exclusion criteria

MGRB participants are consented through the biobank programs of two contributing studies, following protocols previously described [15–17]. Each sample will be from an individual aged 60 years or older with a mean age 75 years across 4,000 participants. Each MGRB participant will have no reported history, or current diagnosis of cardiovascular disease, dementia, or cancer at time of enrolment.

Beyond MGRB inclusion criteria, each sample from the ASPREE study was from a participant aged 75 years or older at time of enrolment, with no reported history of any cancer type. Each sample from the ASPREE study also met the following criteria at time of study enrolment; no clinical diagnosis of atrial fibrillation; no serious illness likely to cause death within the next 5 years (as assessed by general practitioner); no current or recurrent condition with a high risk of major bleeding; no anaemia (haemoglobin > 12 g/dl males, > 11 g/dl females); no current continuous use of other antiplatelet drug or anticoagulant; no systolic blood pressure ≥180 mm Hg and/or a diastolic blood pressure ≥105 mm Hg; no history of dementia or a Modified Mini-Mental State Examination (3MS) score ≤77 [18]; no severe difficulty or an inability to perform any one of the 6 Katz activities of daily living (ADLS)[19].

Beyond MGRB inclusion criteria, each sample from the 45 and Up study also met the following criteria; no record of cancer diagnosis in the NSW Central Cancer Registry; no record of cancer diagnosis in the NSW Admitted Patient Data Collection.

### Phenotypic information

The following data fields will be made available for all MGRB samples through open-access; year of birth, gender, height, weight. For samples from the ASPREE study, waist circumference, blood pressure, fasting blood glucose and status of age-related macular degeneration (AMD) will also be made available through open-access.

## Data Generation

### Library preparation, DNA sequencing, alignment and processing

Whole genome sequencing of MGRB samples will be performed using Illumina HiSeq X sequencers at the Kinghorn Centre for Clinical Genomics (KCCG) under clinically accredited conditions (ISO 15189). Paired-end Illumina TruSeq DNA Nano libraries will be sequenced to one lane per sample. DNA sequences will be mapped to Build 37 of the human reference genome and processed following the Genome Analysis Toolkit (GATK) best practices [20]. Indel realignment and base quality score recalibration of mapped reads will be performed using GATK and best practices parameters; unmapped reads to be left unmodified. GATK HaplotypeCaller will be used to generate g.vcfs from all single-lane realigned and recalibrated BAMs using recommended parameters. All of the raw data will be processed through the Genome One Discovery pipeline [21]. Data will be then analysed using the Hail open-source framework for scalable genetic analysis [22].

### Phased Data Release Plan

MGRB data will be released in three phases (see Table 2). Summary variant frequency data for the MGRB cohort will be made available at the MGRB web portal: https://sgc.garvan.org.au. Complete genotype, phenotype, and raw data are available to potential collaborators upon application. Completion of each phase of sequencing will be followed by a public release of allele frequency data, including an update to the MGRB database, website, portal and beacon.

**Table 2:**
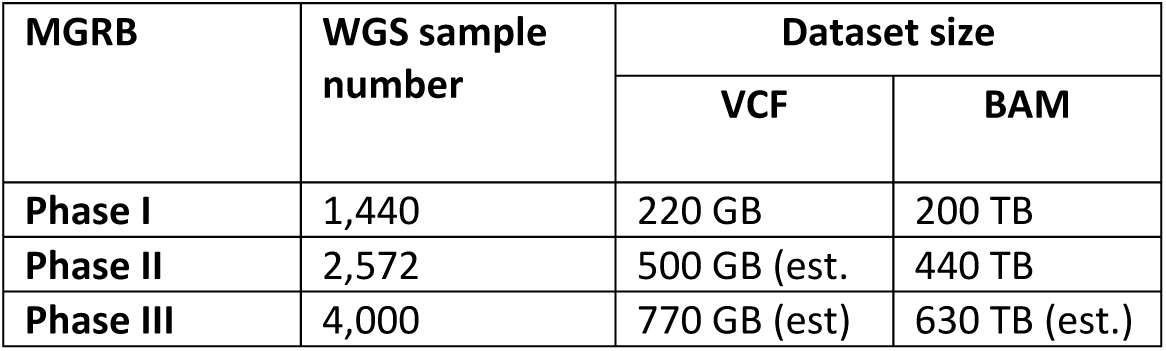
MGRB 3-phase whole-genome sequencing and data release plan

## Data Access

The MGRB Data Access Policy (DAP) summarises the governance applied to individual and institutional access. Curated data will be openly accessible to the international research community through the MGRB website. Preliminary features will include a Beacon, as defined by the Global Alliance for Genomics and Health [23], extensive variant annotation, complex queries (including genetic annotations, and genomic regions), visualisation of variant data (e.g. genome viewer/gene networks) and ultimately, analysis tools for assessing the genetic burden of individual variants and variant subsets.

While basic demographic and phenotypic information will be incorporated into the MGRB data portal, researchers are invited to apply for access to comprehensive genotypic and clinical information to support high-level integrative analysis. To maintain participant privacy and confidentiality, whilst maximising MGRB utility, we have deployed a tiered data management system that determines the richness of data that is made available to researchers (as summarised in the schematic below). This consists of 3 access tiers; Open access, Controlled access and Restricted access.

**Figure 2.**
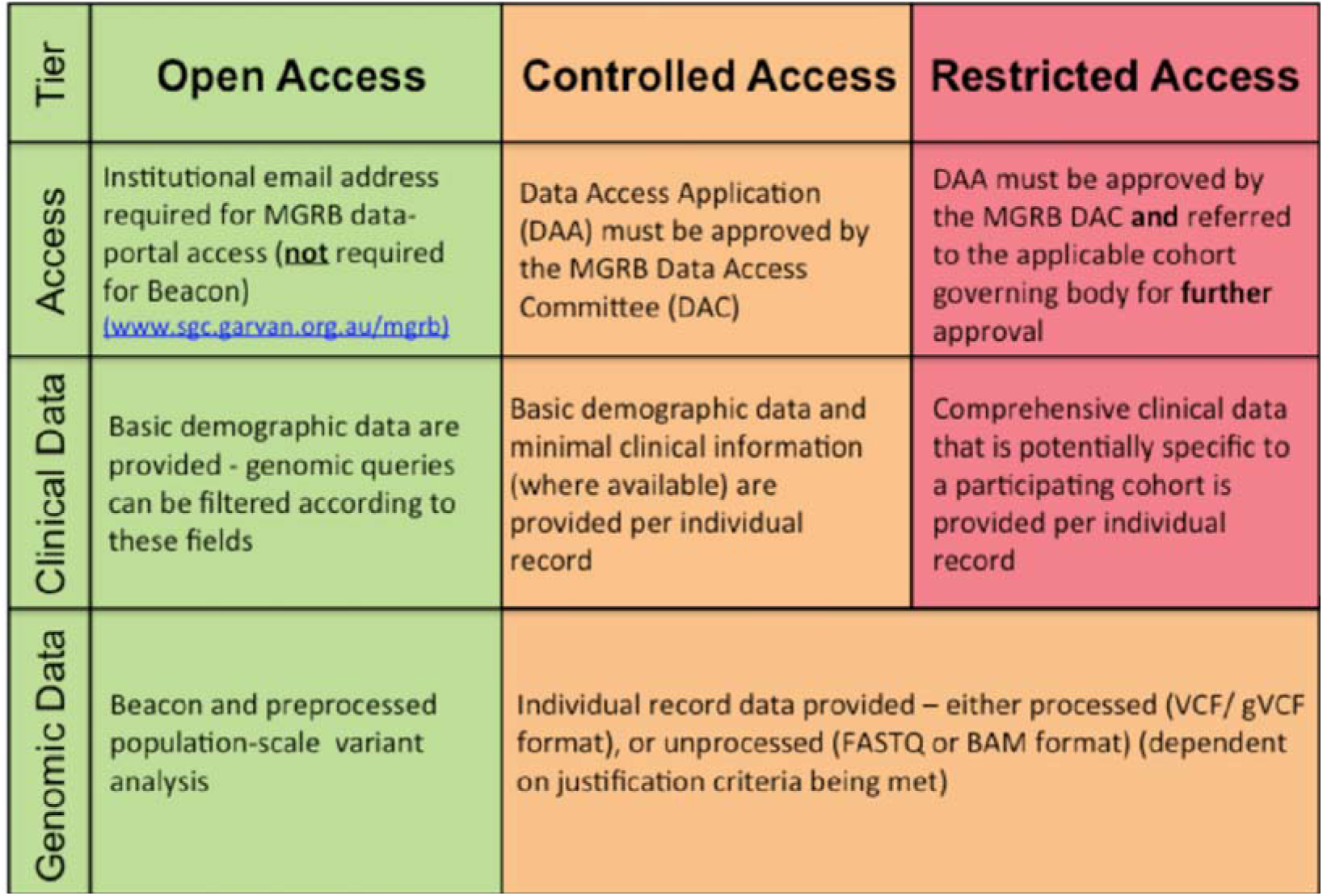
MGRB 3-Tiered Data Access Policy.

The restricted access tier (Tier 3) would involve access to more detailed phenotype and/or clinical information and would require an application, project approval and ethical approval from the ASPREE Presentations, Publications and Ancillary Studies Committee (PPA) or 45andUp Data Access Committee. Notwithstanding internal priorities, and subject to collaborative agreement, both studies commit to fair and reasonable consideration of applications to provide access to restricted access tier data. Individual de-identified processed genomic data would be available to download from the MGRB after execution of a Data Transfer Agreement.

### Open-Source Analytical Framework (*Vectis*)

*Vectis* (a lever in Latin) is a custom-build software environment and collection of modules for the MGRB to support diverse users including clinicians, patients, bench scientists as well as bioinformaticians for the analysis of patient cohorts, of any size, comprising whole genomes, exomes or gene panels. The *Vectis* modules are described in Figure 3 and Table 3.

**Table 3:** Features of the Vectis Open-Source Analytical Framework

| Feature | Description |
| --- | --- |
| Secure login | Two-factor authentication |
| Search | Querying of cohorts using chromosomal coordinates and gene annotations |
| Beacon | Integrated with the Global Alliance for Genomics and Health Beacon Network[20] |
| Explore function | Highly interactive, low latency exploration of cohorts. Explore currently supports the querying of 84 million variants in real-time |
| Interactive graphics | Including lollipop plots of allelic frequencies as well as gene transcripts |
| Variant annotations | Including links out to the original supporting evidence |
| Scalable Variant Store | Enables authorised users to subset patients based on clinical attributes and query actual genotypes at the individual patient level |

**Figure 3.**
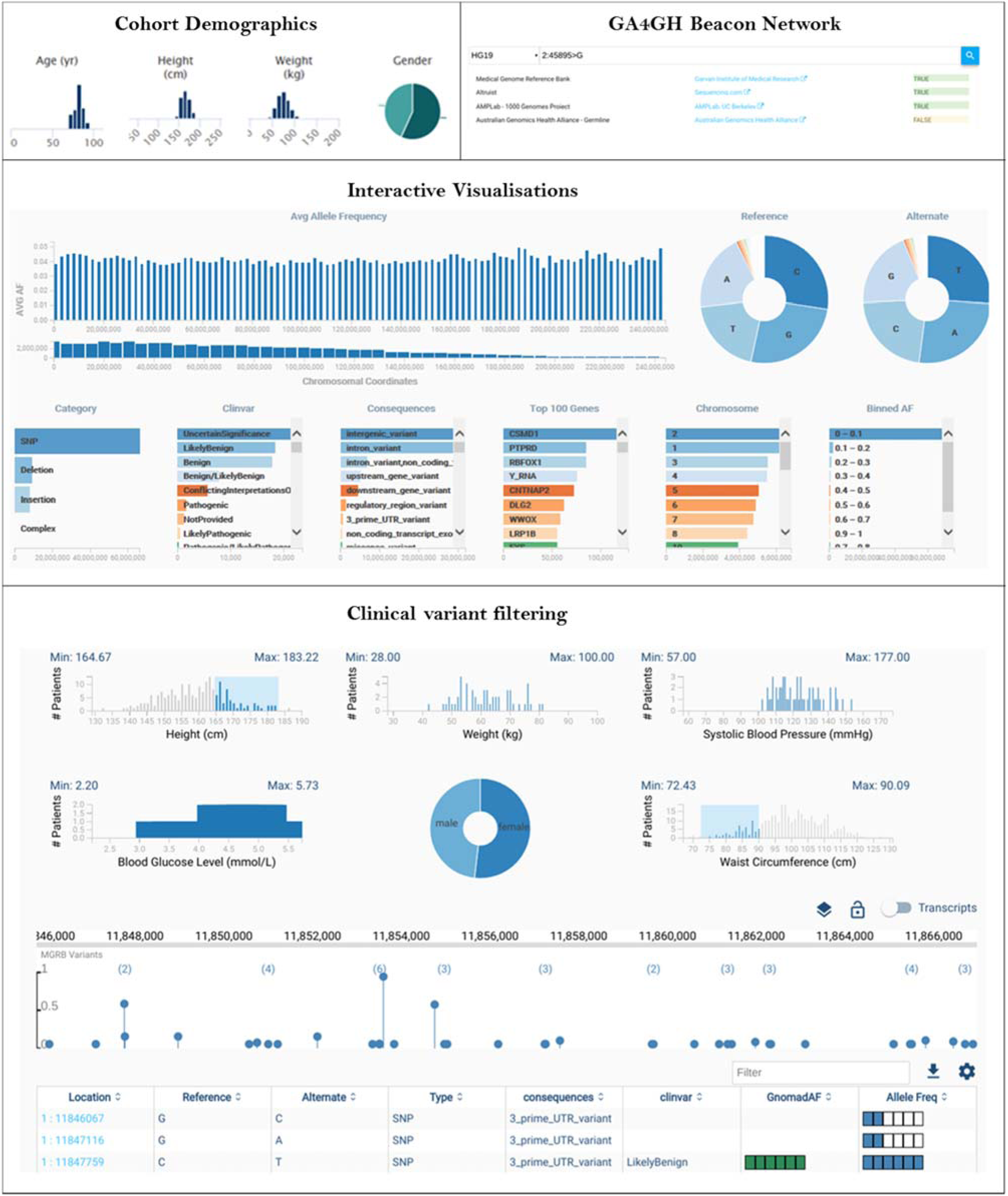
MGRB database functionality and *Vectis* platform.

## Discussion

### Analysis Aims

The overarching aim of the MGRB is to create a catalogue of genome-wide genetic variation in healthy, older individuals, and make data publicly accessible for the clinical genetics and research community. Secondary analytical aims of the MGRB project include, but are not limited to; detection of different variant classes such as single nucleotide variants (SNVs), insertions-deletions (Indels), structural variants (SVs) and copy number changes (CNVs) across the population; clustering of the cohort by ethnicity and other phenotypic factors; examining the frequency and type of clinically significant rare pathogenic variation, in relation to phenotypes; calculating polygenic risk scores (PRS) for a range of conditions, and comparing these scores against population-based and disease-based cohorts; measuring non-germline variation such as telomere length, mtDNA variation and somatic changes in blood in relation to genomic ageing.

### Potential limitations and confounding factors

The MGRB study is limited by a number of factors. The size of the cohort is limited to 4,000 individuals, meaning very rare genetic variants, many of which may be of clinical or biological interest, may not be detected in the dataset by chance, or represented only as singletons (1/4000 or MAF 0.00025). This limit of detection may restrict the sensitivity of MGRB for some applications, compared to larger datasets such as ExAC and Gnomad [4]. However 4,000 samples will still provide a limit of detection of MAF 0.00025, which is below the threshold often used by many studies to define rare variants (MAF<0.01 to MAF<0.001). A related issue is that this Australian cohort is dominated by Caucasian European ancestry, which limits its utility as a variant filter to matched disease populations.

A biological limitation will be the variable penetrance of rare pathogenic variants, even in an elderly population [24,25]. An important consideration is that although the MGRB cohort is an aged, healthy group, it is still possible that rare clinically significant pathogenic variation will be identified, some of which will not be penetrant. Most single gene predispositions, including familial cancers, are not fully penetrant, meaning that less than 100% of individuals with mutations in such genes ever develop the clinical phenotypes associated with that gene mutation, even in older age [24]. The MGRB will give a unique opportunity to overcome the traditional ascertainment bias [26] of human genetics in this regard. However, the detection of a variant in the MGRB alone does not exclude a pathogenic role, where the variable penetrance could be due to genetic or environmental context [27]. This is likely to be particularly important for assigning causality in common diseases, where polygenic effects are likely important. This caveat is something end-users of the data must keep in mind.

A dataset of whole-genome sequences from 600 individuals aged over 80 years has been published previously by the ‘Wellderly’ study [12]. Individuals in this study had no reported chronic diseases and were not taking chronic medications. There are important differences between the Wellderly dataset and the MGRB. Firstly, the number of samples in the MGRB will be significantly higher at 4,000, adding much-needed power and sensitivity for detecting, and filtering, rare variants. MGRB will have sensitivity of 1/4000 (MAF=0.00025) compared to 1/600 (MAF=0.016). Secondly, significant resources within the MGRB have been allocated to delivering open-access data, analytical frameworks and data-sharing mechanisms, for both whole-genome sequencing and phenotypic data. Thirdly, the capacity to detect and report complex and structural genetic variation more readily. Fourthly, the Wellderly study sequenced DNA using the Complete Genomics platform [28], not Illumina which is the technology used by most whole-genome or whole-exome sequencing of reference populations to date [4,5,7–9,11,13]. There are important technical considerations in the cross-compatibility of whole-exome and whole-genome sequencing data for generating population allele frequencies on different sequencing platforms, or data processed using different bioinformatic pipelines [4]. Encouragingly, a recent study showed high concordance between the latest generations of both these sequencing platforms [29].

## Implications

The MGRB has the potential to add another important data resource to the public domain to aid the clinical genetic and research community in the filtering, annotating and assignment of pathogenicity to genetic variants. The unique aspects of the MGRB will include; 1) focus on the healthy elderly, depleted of typical monogenic disease phenotypes; 2) the age of the cohort, average of 75 years, beyond the age of onset for most monogenetic conditions; 3) the availability and access to individual-level VCF and BAM data; and 4) the opportunity to access high quality, comprehensive, longitudinal clinical and phenotypic information. These factors will ensure the MGRB has a unique place alongside other reference populations.

## Conclusion

The MGRB will be the first catalogue of genome-wide genetic variation across thousands of healthy elderly individuals. This will provide an important dataset, resource and much-needed negative control population for clinical genetics and research.

## Acknowledgements

The MGRB was funded by NSW Office of Health and Medical Research - Sydney Genomics Collaborative grant (2014). Authors would like to acknowledge the ASPREE Healthy Ageing Biobank, ASPREE Investigator Group and ASPREE Collaborating Practitioners listed on www.aspree.org. ASPREE was funded by the National Institute on Aging and the National Cancer Institute at the National Institutes of Health (grant number U01AG029824); the National Health and Medical Research Council of Australia (grant numbers 334047, 1127060); Monash University (Australia). The ASPREE Healthy Ageing Biobank was supported by the Commonwealth Scientific and Industrial Research Organisation (Australia), the Victorian Cancer Agency (Australia) and Monash University (Australia). Authors acknowledge the dedicated and skilled staff in Australia and the U.S. for the conduct of the ASPREE trial, and the ASPREE participants who willingly volunteered. Authors would like to acknowledge the 45 and Up Study, managed by the Sax Institute (www.saxinstitute.org.au) in collaboration with major partner Cancer Council NSW; and partners: the National Heart Foundation of Australia (NSW Division); NSW Ministry of Health; NSW Government Family & Community Services – Ageing, Carers and the Disability Council NSW; and the Australian Red Cross Blood Service. We thank the many thousands of people participating in the 45 and Up Study. Authors would like to thank Margo Barr for her contributions to the MGRB project.

